# Non-canonical Features of Pentatricopeptide Repeat Protein-Facilitated RNA Editing in *Arabidopsis* Chloroplasts

**DOI:** 10.1101/544486

**Authors:** Yueming Kelly Sun, Bernard Gutmann, Ian Small

## Abstract

Cytosine (C) to uracil (U) RNA editing in plant mitochondria and chloroplasts is facilitated by site-specific pentatricopeptide repeat (PPR) editing factors. PPR editing factors contain multiple types of PPR motifs, and PPR motifs of the same type also show sequence variations. Therefore, no PPR motifs are invariant within a PPR protein or between different PPR proteins. This work evaluates the functional diversity of PPR motifs in CHLOROPLAST RNA EDITING FACTOR 3 (CREF3). The results indicate that previously overlooked features of PPR editing factors could also contribute to RNA editing activity. In particular, the N-terminal degenerated PPR motifs and the two L1-type PPR motifs in CREF3 are functionally indispensable. Furthermore, PPR motifs of the same type in CREF3 are not interchangeable. These non-canonical features of CREF3 have important implications on the understanding of PPR-facilitated RNA editing in plant organelles.

## Introduction

C-to-U RNA editing is an important post-transcriptional modification process in plant organelles (Takenaka et al., 2013b). Organellar RNA editing is facilitated by organelle-targeted pentatricopeptide repeat (PPR) editing factors (Barkan and Small, 2014). PPR editing factors are site recognition factors, containing tandem helix-loop-helix PPR motifs that bind to the RNA sequence just 5’ to the edited nucleotide in a one-motif to one-base manner. PPR editing factors belong to the PLS-subfamily of PPR proteins, which typically contain the P1-, L1-, S1-, SS-, P2-, L2- and S2-type PPR motifs and the E1- and E2-type PPR-like motifs (Cheng et al., 2016). The motifs are generally arranged following the pattern of (P1-L1-S1)_n_-P2-L2-S2-E1-E2, sometimes with one or more SS motif(s) inserted in between the P1-L1-S1 triplets. PPR editing factors are also involved in the editing reaction, when a deaminase-like DYW domain is located immediately C-terminal to the E2 motif (Wagoner et al., 2015). In some cases, the DYW domain is supplied *in trans* by other proteins (Boussardon et al., 2012; Andres-Colas et al., 2017; Diaz et al., 2017; Guillaumot et al., 2017). PPR editing factors belong to a larger organellar editosome, where multiple other components have been identified (Sun et al., 2016).

Various studies on PPR proteins have revealed the following two main features of PPR-facilitated organellar RNA editing. The most important feature is the PPR-RNA recognition code (Barkan et al., 2012; Takenaka et al., 2013a; Yagi et al., 2013a). The P- and S-type PPR motifs are RNA-recognising motifs. Strong statistical correlation was identified between amino acids encoded at the first and last positions of each motif (i.e. PPR code) and their aligned RNA bases. The statistical correlation is generally weaker between PPR codes encoded in the L1-, L2-, S2-, E1- and E2-type motifs and aligned RNA bases, and has only been experimentally tested in limited examples (Ruwe et al., 2018). Therefore, it was believed that these motifs do not contribute to PPR-RNA recognition as strongly as the P- and S-type PPR motifs. The correlation between each possible PPR code and RNA base is used to score a PPR motif against its aligned RNA base. Sum of the score for each PPR motif indicates the overall degree of matching between a PPR protein and its RNA target. It means that PPR motifs containing codes that have strong correlation with RNA bases would contribute more to the scoring. Namely, each PPR motif weighs differently based on different degrees of statistical correlation between its PPR code and aligned RNA bases. The other important feature of PPR-facilitated RNA editing is the one-motif to one-base modularity for RNA recognition, where each PPR motif is an independent RNA base-recognising module. It implies the potential for PPR motifs to be shuffled within a single protein and between different proteins that could lead to changes in RNA targeting specificity (Yagi et al., 2014).

Here we use CHLOROPLAST RNA EDITING FACTOR 3 (CREF3, encoded by *AT3G14330*), the site-recognition factor for the *psbE* editing site in *Arabidopsis* chloroplasts (Yagi et al., 2013b), as an example to elucidate features of PPR editing factors not previously described. We show that these non-canonical features could also affect the efficiency of PPR-facilitated organellar RNA editing.

## Results

### An *in vivo* system for evaluating CREF3 variants with RNA editing as reporter

According to the statistical correlation between the fifth and last positions of PPR motifs and aligned RNA bases, a scoring matrix was generated for CREF3 indicating its targeting preferences (Figure 1a). The 5’ *cis*-elements of the *psbE* editing site in *Arabidopsis* chloroplasts were aligned with CREF3 PPR motifs. The HMMER scores of CREF3 PPR motifs are plotted in a bar chart above the schematic illustration of CREF3 motif arrangement, indicating how much a CREF3 motif resemble the typical PPR motif of its type.

**Figure 1.**
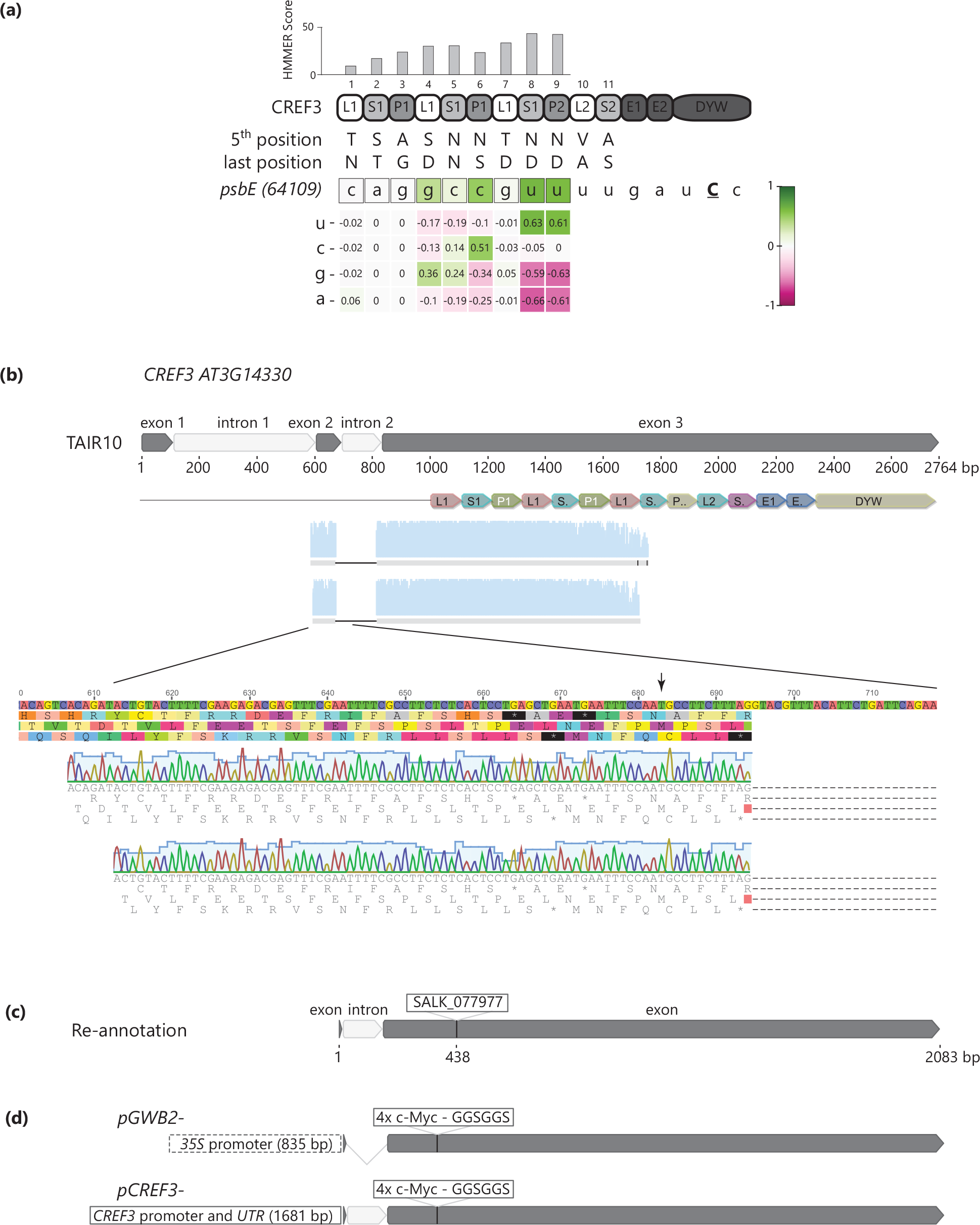
An *in vivo* system for evaluating CREF3 variants with RNA editing as reporter (a) Alignment of CREF3 motifs with the *psbE* editing site in *Arabidopsis* chloroplasts. <u>C</u> indicates the edited cytidine. The nucleotides are coloured according to the alignment scores (Sun et al., 2018). (b) The *CREF3 (AT3G14330)* gene model suggested by the TAIR database (http://www.arabidopsis.org), with two introns in the presequence; And the *CREF3* gene model corrected by 5’RACE, with the start codon shifted 681 bp downstream, and only one intron present in the presequence. (c) Map of the *CREF3* T-DNA insertion in SALK_077977. T-DNA is inserted in from of the nucleotide 1119 in the gene model suggested by TAIR, within the first PPR motif (1-L). (d) Illustration of the *CREF3* transgene models. Top: The expression of *CREF3* or variants thereof is driven by the CaMV *35S* promoter, without its intron, tagged with four copies of c-Myc, followed by a GGSGGS flexible linker in front of the first PPR motif; Bottom: The expression of *CREF3* or variants thereof is driven by the native *CREF3* promoter and *UTR* sequence, with its intron included, also tagged with four copies of c-Myc, followed by a flexible GGSGGS linker in front of the first PPR motif.

In *Arabidopsis*, full-length *CREF3* transcripts could not be amplified according to the gene annotation in TAIR10 (Figure 1b). RNA-seq datasets (Dubreuil et al., 2018) also suggest that there is no reads mapped to the first annotated exon and intron of *CREF3*. Therefore, rapid amplification of 5’ cDNA ends (5’RACE) was performed to map the 5’ end of *CREF3* transcripts. The start codon is re-predicted to be 681 bp downstream from the original annotation in TAIR10, followed by a confirmed intron. The T-DNA insertion in the SALK_077977 line, which represents a *null* mutation of *CREF3* (Yagi et al., 2013b), is mapped upstream of the 438^th^ nucleotide in the re-annotated *CREF3* gene model, within the first PPR motif of 1-L1 (Figure 1c).

To evaluate the function of CREF3 PPR motifs, tagged CREF3 variants modified at the motif(s) of interest were expressed in *cref3* mutant background, and RNA editing at relevant sites was quantified as functional reporter. The CREF3 variants were either expressed from the pGWB2 vector (EMBL), or from the home-made pCREF3 vector, modified from the plant expression vector pAlligator2 (Bensmihen et al., 2004) (Figure 1d). The pGWB2-based *CREF3* constructs are driven by a *CaMV 35S* promoter, with the *CREF3* intron removed, as well as with 4x *c-Myc* tags and a flexible linker (GGSGGS) inserted immediately upstream of the first PPR motif 1-L1. The pCREF3-based *CREF3* constructs contain *CREF3* genomic sequence starting from 1681 bp upstream of the re-predicted start codon. Four copies of the c-Myc tag and a flexible linker (GGSGGS) are inserted in front of the first PPR motif 1-L1. Both the pGWB2-based (Figure 3b) and the pCREF3-based (Figure 2c and 4b) *CREF3* wild type constructs fully complement the *cref3* mutant in terms of *psbE* editing.

**Figure 2.**
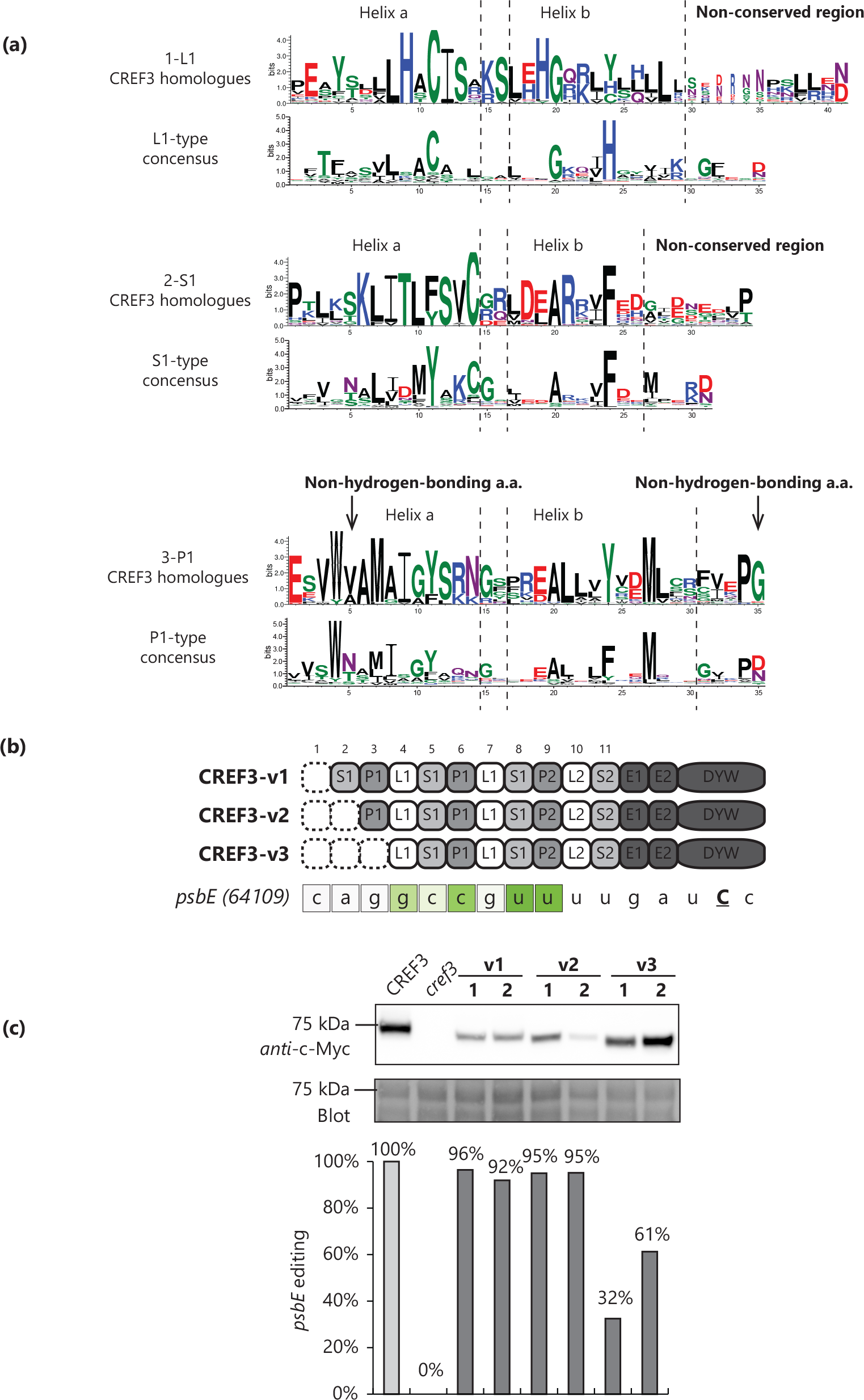
CREF3 N-terminal truncations (a) Alignments of CREF3 motif consensus sequences with the corresponding L, S, or P motif consensus sequences (Cheng et al., 2016). (b) Illustration of CREF3 N-terminal truncation variants. In CREF3-v1, 1-L was truncated; In CREF3-v2, 1-L and 2-S were truncated; In CREF3-v3, 1-L, 2-S and 3-P were truncated. (c) Protein and RNA analyses of transgenic plants expressing CREF3 N-terminal truncation variants in the *cref3* mutant background. Two independent transgenic lines were selected for each CREF3 variant. Top: Immunoblotting with the anti-c-Myc antibody; Middle: Blot image after protein transfer prior to antibody incubation; Bottom: Quantification of editing at the *psbE* site.

**Figure 3.**
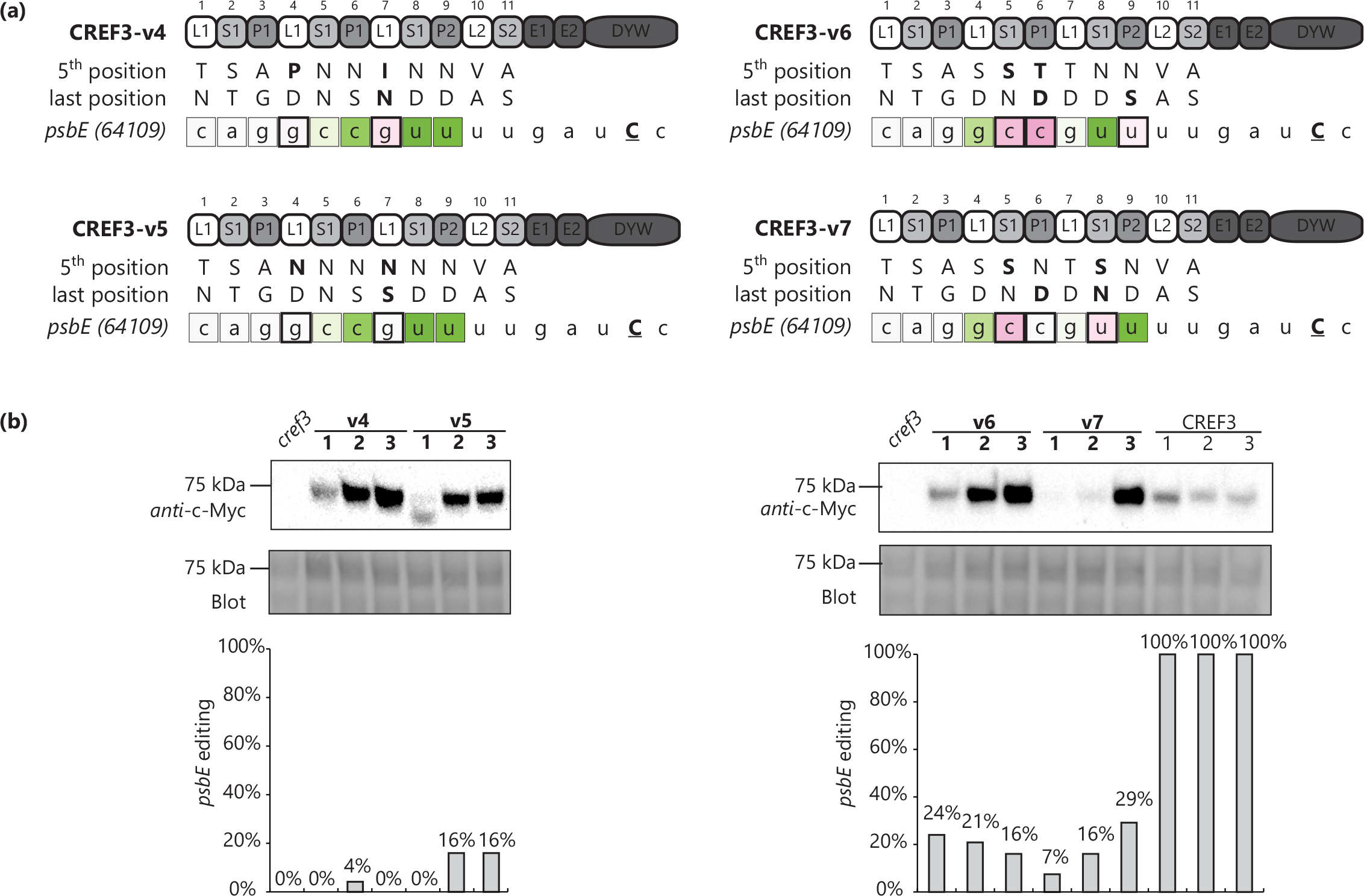
CREF3 variants with modified PPR-RNA code (a) Illustration of CREF3 L motif variants (CREF3-v4 and v5) and P/S motif variants (CREF3-v6 and v7) with modified PPR-RNA code and their new alignments with the *psbE* site. (b) Protein and RNA analyses of transgenic plants expressing CREF3 variants in the *cref3* mutant background. Three independent transgenic lines were selected for each CREF3 variant. Top: Immunoblotting with the anti-c-Myc antibody; Middle: Blot image after protein transfer prior to antibody incubation; Bottom: Quantification of editing at the *psbE* site.

### The degenerate N-terminal motifs of CREF3 are indispensable

The first three PPR motifs at the N-terminus of CREF3 do not show strong statistical correlation with the aligned RNA bases (Figure 1a). Especially, the fifth and last positions of the 3-P1 [AG] motif (the 3^rd^ PPR motif that is of P1-type encoding A and G at the fifth and last position respectively) encoded as alanine (A) and glycine (G) are not capable of forming hydrogen bonds, while hydrogen bonding plays an important part in PPR-RNA recognition (Shen et al., 2016). Moreover, the 1-L1 [TN] and 2-S1 [ST] motifs show lower HMMER scores compared to the other PPR motifs in CREF3 (Figure 1a), indicating that these are not typical PPR motifs.

Sequence logos of CREF3 motifs were generated from 17 CREF3 homologues (Figure 2a). Both the 1-L1 and the 2-S1 motifs contain a non-conserved region following helix b. Within this region, the conserved amino acids in the corresponding consensus sequences are lost. In the 1-L1 motif, the conserved G31 and F32 amino acids are lost. In the 2-S1 motif, the conserved M27 and R/K30 amino acids are lost. Moreover, the last position of 2-S1 no longer encodes a typical RNA-recognising amino acid D or N, instead, P or T is encoded. The fifth and last positions of the 3-P1 motif consistently encode non-hydrogen-bonding amino acids V/A and G. Therefore, it was hypothesised that the motifs 1-L1, 2-S1 and 3-P1 do not function as canonical RNA-recognising motifs and that they may be dispensable in CREF3.

Serial truncation of the CREF3 N-terminal motifs was performed, and three variants were generated - CREF3-v1 with the 1-L1 motif truncated, CREF3-v2 with the 1-L1 and 2-S1 motifs truncated, and CREF3-v3 with all three motifs truncated (Figure 2b). As shown in Figure 2c, all three variants were expressed and accumulated in the *cref3* mutant background. CREF3-v1 and v2 complemented the *psbE* editing phenotype similar to the wild type level, whereas CREF3-v3 only partially complemented *psbE* editing to 30%-60%, correlating with the protein expression level. These results indicate that the 1-L1 and 2-S1 motifs may be dispensable, however, the 3-P1 motif is required for optimal editing activity of CREF3.

### Two L1-type motifs of CREF3 are critically involved in RNA recognition

Previously, robust correlation between L1-type motifs and aligned RNA bases could not be established with bioinformatics (Barkan et al., 2012; Takenaka et al., 2013a; Yagi et al., 2013a). The correlation between canonical codes and aligned RNA bases are either weak (S_5_N_last_-A and S_5_D_last_-G) or non-existent (N_5_S_last_-C, N_5_D_last_-U). Besides, there is correlation between non-canonical codes and aligned RNA bases (e.g. P_5_D_last_-U, I_5_N_last_-C/U), while only one amino acid position may be capable of forming hydrogen bonds. Therefore, L1-type motifs are generally believed not to be important in RNA recognition. The crystal structure of a consensus-based synthetic PLS protein, and the RNA binding assays conducted with the same design, showed that the synthetic L motifs can recognise RNA yet only upon binding by MORF9 protein (Yan et al., 2017), which is a critical component identified in the editosome (Takenaka et al., 2012).

CREF3 presents a special case, where the fifth and last positions of 4-L1 [SD] and 7-L1 [TD] motifs encode canonical amino acid combinations matching the aligned RNA bases (G in both cases). In addition, these two positions show specificity towards G in an *in vitro* editing assay (Hayes and Hanson, 2007). Furthermore, these two positions are the only positions that differentiate between the *psbE* and *petL* editing sites in *Arabidopsis* chloroplast, with *psbE* edited by CREF3, and *petL* apparently not edited by CREF3. Therefore, it is hypothesised that the 4-L1 and 7-L1 motifs in CREF3 are involved in RNA recognition. Since *psbE* editing is not defective in *morf9* mutants (Takenaka et al., 2012; Bentolila et al., 2013), the function of CREF3-L1 motifs is MORF9 independent.

Two L1 motif variants were generated with modifications at the fifth and last positions of 4-L1 and 7-L1, aiming to switch the targeting specificity from G (a purine) to C or U (pyrimidines). Two sets of codes were considered – 1) the L1-specific codes PD-U and IN-C/U (Figure 3a, v4); and 2) the canonical codes NS-C and ND-U (Figure 3a, v5). In comparison, another two CREF3 variants were generated with modifications at the fifth and last positions of P1- or S1-type motifs (Figure 3a, v6 and v7). As shown in Figure 3b, all four CREF3 variants were expressed and accumulated in the transgenic plants, however, with *psbE* editing dramatically compromised compared to the plants expressing the wild type CREF3. There is no new editing events detected by RNA-seq in any of these variants. Therefore, the 4-L1 and 7-L1 motifs in CREF3 are involved in PPR-RNA interaction, and their fifth and last positions may be involved in G recognition in a similar manner compared to the P- and S-type motifs. Moreover, it appears that each motif contributes differently to CREF3-facilitated *psbE* editing. Modifications of two L1-type motifs led to similar, if not more dramatic, effects on RNA editing compared to modifications of three P1- or S1-type motifs. It indicates that the contribution of each PPR motif to RNA editing is not necessarily determined by the statistical correlation between the PPR code and aligned RNA base.

### Two similar P1-L1-S1 triplets in CREF3 are not readily interchangeable

It is questionable whether PPR motifs of the same type are interchangeable. The six RNA-recognising motifs in CREF3 can be split into two LSP triplets recognising similar nucleotide combinations “GYY” (Y=C or U) at the *psbE* editing site (Figure 4a). The triplet A consists of motifs 4 to 6 (L1-S1-P1). The triplet B consists of motifs 7 to 9 (L1-S1-P2). The P1/P2 classification on 6-P1 and 9-P2 is due to their positions in CREF3, rather than their motif sequences (Cheng et al., 2016). It is hypothesised that these two LSP triplets are functionally equivalent in CREF3. To test the hypothesis, two CREF3 variants were generated by replacing the triplet B with A or A with B, and recoding to maintain the original PPR code aligned to each RNA base (Figure 4a, v8 and v9).

**Figure 4.**
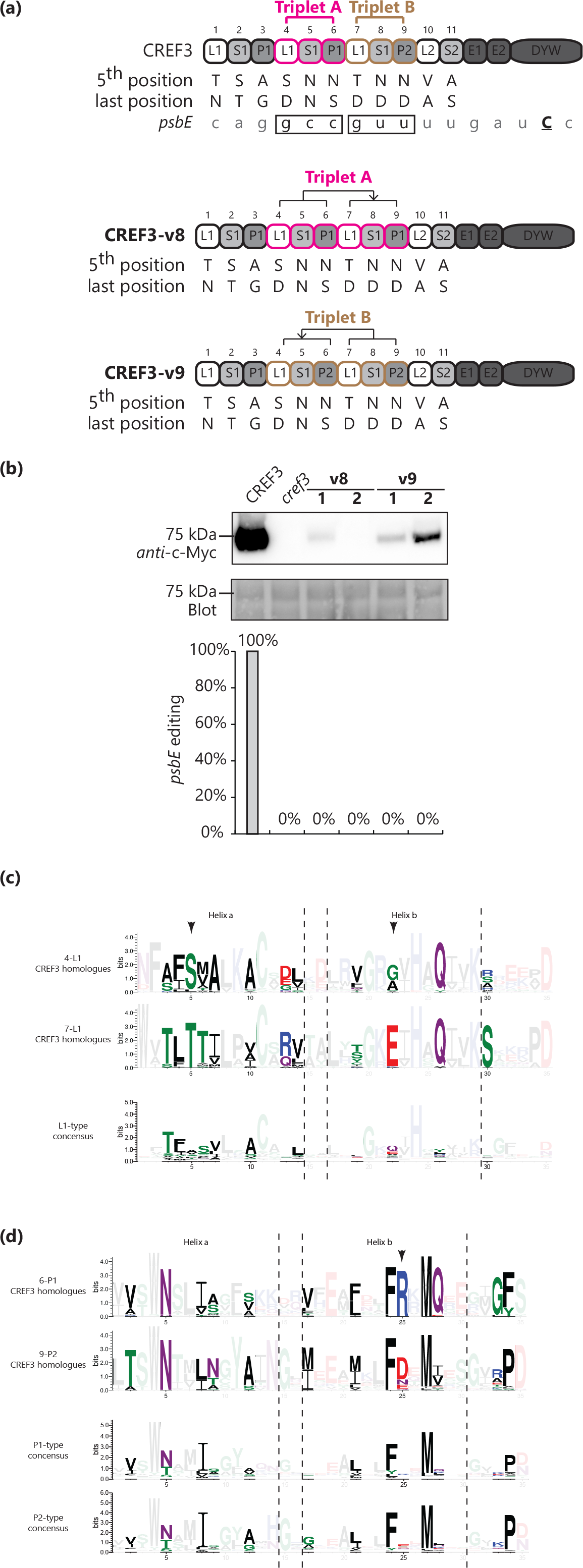
Swapping the motif triplets in CREF3 (a) Illustration of two sets of CREF3 LSP motif triplets both targeting the nucleotides “GYY”. Triplet A: motifs 4-6; Triplet B: motifs 7-9. Illustration of CREF3 triplet swapping variants. In CREF3-v8, the backbone of triplet B was replaced by triplet A while maintaining the matches with the *psbE* editing site through the fifth and last amino acids of each motif. In CREF3-v9, the backbone of triplet A was replaced by triplet B while maintaining the matches with the *psbE* editing site. (b) Protein and RNA analyses of transgenic plants expressing CREF3-v8 or CREF3-v9 in the *cref3* mutant background. Two independent transgenic lines were selected for each CREF3 variant. Top: Immunoblotting with the anti-c-Myc antibody; Middle: Blot image after protein transfer prior to antibody incubation; Bottom: Quantification of editing at the *psbE* site. (c) Alignments of CREF3 motif consensus sequences with the corresponding L motif consensus sequence (Cheng et al., 2016). Predicted contact points with the adjacent P motif are highlighted. (d) Alignments of CREF3 motif consensus sequences with the corresponding P motif consensus sequences (Cheng et al., 2016). Predicted contact points with the adjacent L motif are highlighted.

As shown in Figure 4b, CREF3-v8 and v9 were expressed not nearly as well as the wild type CREF3, and neither of them complemented *psbE* editing. The P1-L1-S1 triplets placed at non-native positions could destabilise CREF3 despite being the same type. The results indicate that the two P1-L1-S1 triplets within CREF3 are not functionally equivalent.

Since CREF3 motif swaps alter neighbouring PPR motif pairs (P1-L1), the structural instability may be due to incompatible amino acids at the motif interfaces. Sequence logos of 4-L1, 7-L1, 6-P1 and 9-P2 were generated from 17 CREF3 homologues and aligned with the sequence logos of the corresponding L1-, P1- and P2-type motifs (Figure 4c and 4d). Predicted amino acid positions in contact with the neighbouring motif (Cheng et al., 2016) are highlighted. As shown in Figure 4c, the positions in L1-type motif in contact with the previous P1-type motif are mostly non-conserved between L1-type motifs. However, some of these positions become conserved within the 4-L1 or 7-L1 homologue groups. Moreover, these positions show different conservation patterns between the 4-L1 and 7-L1 groups. As shown in Figure 4d, the positions in P-type motifs that are in contact with the following L-type motifs are similar between the P1- and P2-type motifs. Some of these positions also diverge between CREF3 6-P1 and 9-P2 homologue groups. The above sequence comparison implies that each of the 4-L1 and the 7-L1 motifs, and of the 6-P1 and 9-P2 motifs, are compatible with different neighbouring motifs. As a result, CREF3-v8 and v9 variants with altered P1-L1 motif junctions are not stable.

## Discussion

### Disagreement between statistical and observed contribution of a PPR motif to RNA editing

In contradiction to the previous conclusions (Barkan et al., 2012; Takenaka et al., 2013a; Yagi et al., 2013a; Yan et al., 2017), two L1-type motifs in CREF3 contribute strongly to *psbE* editing, following the canonical PPR recognition code, and independent of MORF9. Surprisingly, these two L1 motifs contribute as strongly to the overall editing activity of CREF3, as compared to the P- and S-type motifs that show stronger statistical correlation with the aligned RNA bases. The disagreement between statistical and observed correlations is consistent with the previously tested example of CLB19 (Kindgren et al., 2015), where the 1-P1 [TN] and 3-S1 [TN] motifs do not appear to affect the RNA-recognition specificity, despite showing strong statistical correlation with the aligned RNA bases (A in both cases).

The intensity of chemical interaction between a PPR motif and its aligned RNA base may explain the observation of differential contribution (Shen et al., 2016). For example, one could argue that the L1 motifs in CREF3 contribute more to RNA recognition because they interact with the RNA base G through three hydrogen bonds, whereas the P- or S-type motifs in CREF3 interact with RNA bases C or U via two hydrogen bonds. However, this explanation cannot account for all the observations. For example, it does not explain why the A-aligning motifs (1-P1 [TN] and 3-S1 [TN]) contribute less compared to the C-aligning motif (8-P2 [ND]) in CLB19 (Kindgren et al., 2015).

Taken together, PPR-RNA editing specificity is not solely determined by the statistics-based canonical PPR-RNA code focusing on the P- or S-type motifs. Contribution-weighing factors for each PPR motif need to be determined for more accurate prediction of editing sites in the future.

### PPR proteins are organised in a coherent manner

This work demonstrates that it is not straightforward to swap CREF3 PPR motifs with others of the same type due to protein stability issues. This observation reveals the complication behind the modularity concept of PPR-RNA recognition, and highlights the requirement for compatibility at the PPR motif interface. Modelling (Cheng et al., 2016) and structures (Yan et al., 2017) of PLS proteins suggest substantial interactions between PPR motifs, which are primarily van der Waals interactions. In general, the interaction between each pair of adjacent PPR motifs could also contribute to the overall conformational plasticity of a PPR protein (Shen et al., 2016), which ultimately controls the compression of the superhelix upon RNA binding. Therefore, any subtle differences introduced to the interaction between PPR motifs may accumulate and affect the overall superhelical architecture of a PPR protein and its RNA binding capacity.

Predicted contact positions with neighbouring PPR motifs are mostly variable between PPR motifs of the same type (Cheng et al., 2016), however, are conserved between the homologous motifs in CREF3. The divergent evolution of these contact positions implies that an overall coherent organisation of PPR motifs may be evolutionarily conserved.

### Relationship between PPR sequence-degeneracy and function-degeneracy

The N-terminal PPR motifs of CREF3 contribute to RNA editing despite the degeneracy in their sequences. According to *in vitro* RNA binding assays previously conducted with the PPR editing factors CRR21, OTP80, CRR4, CRR28 and OTP85 (Okuda et al., 2014), the interaction between the first two PPR motifs and the aligned RNA bases contribute to the overall binding affinity. In retrospect, we found that these motifs also have low HMMER scores and/or encode non-hydrogen-bonding amino acids at the fifth and last position: CRR21-1-P1 [FG] (HMMER score 0), OTP80-2-S1 [FN] (HMMER score 4.2), CRR4-2-S1 [VD] (HMMER score -1), CRR28-1-S1 (HMMER score 8.2) and OTP85-1-SS [ID] (HMMER score 9.2). These evidence support our conclusion that PPR sequence-degeneracy at the N-termini does not necessarily indicate function-degeneracy, and imply that these motifs may contribute to RNA binding affinity in a non-specific manner.

In general, it is not uncommon that repeat proteins carry sequence degeneracy that is also functional at their N-termini. For example, the DNA-recognising TALE (transcription activator-like effectors) proteins contain four degenerate repeats at the N-terminal region (NTR). It was shown that the NTR is crucial to the activity of TALE fusion proteins *in vivo* (Mussolino et al., 2011) by interacting non-specifically with dsDNA (Gao et al., 2012) and facilitating one-dimensional scanning along the DNA candidates (Cuculis et al., 2015).

Taken together, the non-canonical features of PPR-facilitated RNA editing demonstrated in this work create new avenues for investigating the mechanism of this important post-transcriptional processing events in organelles, which could eventually guide protein engineering efforts for developing programmable RNA editing tools *in planta*.

## Methods

### Rapid amplification of 5’ cDNA ends (5’RACE)

Rapid amplification of 5’ cDNA ends (5’RACE) of *CREF3* transcripts was performed using SMARTer RACE 5’/3’ Kit (Clontech) according to the manufacturer’s instructions. 900 ng DNase-treated Col-0 RNA was used for cDNA synthesis. CREF3-specific fragments were amplified from undiluted cDNA using PrimeSTAR polymerase according to the manufacturer’s instructions, with the following primer combination.

Forward: 10×UPM provided by the SMARTer Kit

Reverse: ACCAGCTCGCCCCAAGATGTCAACCAAACAAGCATAATGC (*CREF3* gene-specific).

The PCR products were gel purified, cloned into pGEMT-Easy (Promega) according to the manufacturer’s instructions, and sequenced.

### Cloning of plant transformation constructs

For cloning into the plant expression vector pGWB2 (EMBL), the *CREF3* gene or variants thereof were mutagenised as needed, then assembled and amplified with *attB* recombination sites using PrimeSTAR polymerase (Clontech). The PCR product was purified by either QIAquick Gel Extraction Kit or QIAquick PCR Purification Kit (Qiagen), cloned into the donor vector pDONR207 using Gateway BP Clonase (Invitrogen), and transformed into competent *E. coli* cells (DH5α). The *CREF3* gene or variants thereof were then transferred from the donor vector pDONR207 to the plant expression vector pGWB2 (EMBL) using Gateway LR Clonase (Invitrogen), and transformed into competent *E. coli* cells (DH5α). Positive clones for each construct were confirmed by Sanger sequencing. The verified constructs were transformed into competent *Agrobacterium tumefaciens* cells (GV3101).

For construction of the plant expression vector pAEF3-Ali, three fragments were prepared for Gibson assembly: 1) the plant expression vector pAlligator 2 (Bensmihen et al., 2004) was digested with *Hin*dIII and *Sal*I to remove the CaMV *35S* promoter and the Gateway cassette; 2) the native *CREF3* promoter and UTR sequence was amplified with appropriate overlapping arms from Col-0 genomic DNA; 3) the 4×c-Myc tag was amplified with the GGSGGS linker, the *Asc*I restriction site, and appropriate overlapping arms from the plant expression vector pGWB17 (EMBL). About 50 fmol of each fragment were combined in the Gibson assembly reaction, incubated at 50°C for 60 min, and transformed into competent *E. coli* cells (DH5α). Positive clones were confirmed by Sanger sequencing.

For cloning into the plant expression vector pAEF3-Ali, the following fragments were prepared for Gibson assembly: 1) pAEF3-Ali was linearised with *Asc*I; 2) one or more fragments of the *CREF3* gene or variants thereof were mutagenised as needed and amplified with appropriate overlapping arms using PrimeSTAR polymerase (Clontech). Each fragment was combined in the Gibson assembly reaction according to the following guidelines – 25 fmol of fragments larger than 1 kb; 75 fmol for fragments between 500 bp and 1kb; and 125 fmol for fragments smaller than 500 bp. The Gibson assembly reaction was incubated at 50°C for 1-12 hrs depending on the number of fragments to be assembled, and transformed into competent *E. coli* cells (DH5α). Positive clones for each construct were confirmed by Sanger sequencing.

### Plant growth, transformation, and selection

Arabidopsis seeds harvested from homozygous *cref3* mutant plants (SALK_077977) were surface sterilised with 70% Ethanol + 0.05% Triton-X100 for 5 min and washed with 100% ethanol before being dried in a fume hood. Sterilised seeds were sowed on plates (half-strength MS medium and 0.8% agar), stratified at 4 ℃ in the dark for 3 days, germinated and grown under long-day conditions (16h light/8h dark cycle, approximately 120 μmol photons m^−2^ s^−1^). The primary stems were trimmed to induce branching. Upon flowering, the *cref3* plants were transformed with constructs of *CREF3* or its variants by floral dip, a method for *Agrobacterium*-mediated transformation of Arabidopsis (Clough and Bent, 1998).

Seeds harvested from plants dipped with the pGWB2 constructs were selected on half-strength MS agar plates with Hygromycin B (25 μg/ml). Surviving primary transformants (T_1_) were transferred to soil and screened for transgene expression by immunoblotting against the c-Myc tag.

Seeds harvested from plants dipped with the pAEF3-Ali constructs were sowed on half-strength MS agar and selected under a stereo microscope fitted with a fluorescence adapter (Royal Blue, Nightsea) to detect GFP expression in the seed coat. Glowing seeds of the primary transformants (T_1_) were transferred to a new plate, grown for about 10 days, then transferred to soil and screened for transgene expression by immunoblotting against the c-Myc tag.

Wild type Col-0 plants were grown and transformed using the same protocols as *cref3*. The chloroplast editing factor mutants, *flv* (SALK_139995) and *ys1* (SALK_123515), were grown and transformed using similar protocols, except that 0.5% sucrose was added to the half-strength MS medium.

### Protein extraction and immunoblotting

Three leaves from a 3-week old plant were collected into a 2-ml tube carrying one stainless steel ball (3-mm diameter) and the tube was snap-frozen in liquid nitrogen. The frozen tissue samples were held in a rack pre-chilled in liquid nitrogen and ground to a fine powder using a mixer mill at 30/s for 1 min. Soluble proteins were extracted using 50 μl of protein extraction buffer (50 mM Tris-HCl pH 7.5, 150 mM NaCl, 10% glycerol, 0.1% Tween-20, 1 mM DTT and 1×Complete protease inhibitors [Roche]), incubated on ice for 5 min. Tissue debris was pelleted by centrifugation at 20,817 *g*, 4°C, for 8 min. 20 μl of soluble proteins were combined with 4 μl of 6×SDS-PAGE protein sample buffer, denatured at 95°C for 5 min, and separated on a 10% SDS-PAGE gel (TGX Stain-Free FastCast, Bio-rad). The protein gel was directly imaged (Gel Doc, Bio-rad), and blotted onto PVDF membrane (Immun-Blot, Bio-rad) using a semi-dry transfer cell (Trans-Blot, Bio-rad). The Blot was directly imaged after protein transfer, blocked in 1×TBS-T with 1% Blocking Reagent (Roche) at room temperature for 1 h, and incubated with anti-c-Myc primary antibody solution (GenScript, 1: 2,000 dilution in 1×TBS-T with 0.2% Blocking Reagent [Roche]) overnight at 4°C. After removing the primary antibody solution, the blot was washed with 1×TBS-T solution (4×5min) and incubated with anti-Mouse IgG-HRP secondary antibody solution (Sigma, 1:10,000 dilution in 1×TBS-T with 0.2% Blocking Reagent [Roche]) for 1-2 hrs. After removing the secondary antibody solution, blots were washed with 1×TBS-T solution (4×5min). HRP (horseradish peroxidase) activity was detected using Clarity Western ECL substrate (Bio-rad). Chemiluminescence images were collected with an ImageQuantRT ECL system (GE Healthcare).

### RNA extraction and editing analysis

Total RNA from seedling or leaf tissue of the primary transformants (T_1_) was isolated using the PureZOL reagent (Bio-Rad) according to the manufacturer’s instructions. Total RNA was treated with TURBO DNase (Ambion) according to the manufacturer’s instructions. Completion of DNase treatment was verified by PCR targeting chloroplast genomic DNA. Complementary DNA (cDNA) was synthesised from the DNase-treated RNA using random primers and SuperScript III reverse transcriptase (Invitrogen) according to the manufacturer’s instructions. Synthesised cDNA was diluted and used as PCR template. PCR was conducted using PrimeSTAR polymerase (Clontech) according to the manufacturer’s instructions. The following primer pairs and conditions were used.

For *psbE_64109* only (80 bp):

Forward: AAGGCATTCCATTAATAACAGG
Reverse: TGGGTCCTCCTAAAAAGATCTAC
40 cycles of 98°C, 10 sec; 60°C, 15 sec; 72°C, 5 sec

For both *psbE_64109* and *psbE_64078* (366 bp):

Forward: ACAGGAGAACGTTCTTTTGC
Reverse: TCGTTGGATGAACTGCATTG
40 cycles of 98°C, 10 sec; 60°C, 15 sec; 72°C, 20 sec

For *ndhB_95252* (99 bp):

Forward: GGCTCTCTCTTTAGCTCTATGTC
Reverse: GCCTGCCATCCACACCAGAATA
40 cycles of 98°C, 10 sec; 60°C, 15 sec; 72°C, 5 sec

For *ycf1_128321* (276 bp):

Forward: GGACCAAGAGGTATCCACCGA
Reverse: ACGAGAGTTACAAATGGTTTTTCAAACC
40 cycles of 98°C, 10 sec; 58°C, 15 sec; 72°C, 15 sec

For *rpoC1_21806* (324 bp):

Forward: TTTTCTTTTGCTAGGCCCATAACT
Reverse: CTTAGCTAATTCCATACGTCTAAC
40 cycles of 98°C, 10 sec; 60°C, 15 sec; 72°C, 20 sec

For *rpoB_25992* (318 bp):

Forward: TTTGGAAAACCAGTAGGAATATGC
Reverse: CTCGTAGATTCAAACCCATAGC
40 cycles of 98°C, 10 sec; 60°C, 15 sec; 72°C, 20 sec

RNA editing was quantified either by poisoned primer extension (PPE) as described by Chateigner-Boutin and Small (2007) or by Sanger sequencing of the PCR products. The following primers and dideoxynucleotides were used for PPE.

For *psbE_64109*, 6’FAM-CTAAATTCATCGAGTTGTTCCAAAG, ddG

For *psbE_64078*, 6’FAM-TTTGGAACAACTCGATGAATTTAGTAGAT, ddC

For *ndhB_95252*, 6’FAM-CTATGTCTCTTATCCCTAGGAGGTCTTCCT, ddT

For *ycf1_128321*, 6’FAM-AGTTTTATAGTTATAGTATGTTCGAACGTG, ddG

### DNA extraction and genotyping

For less than 12 samples, DNA was extracted using the DNeasy Plant Kit (Qiagen). For more than 12 samples, DNA was extracted using a high-throughput method. Briefly, plant tissues were harvested in strip cluster tubes each carrying one stainless steel ball (3-mm diameter) and ground in 300 μl DNA extraction buffer (100 mM Tris-HCl pH 8.0, 50 mM EDTA pH 8.0, 500 mM NaCl, and 1% PVP-40). Proteins were precipitated by adding 37.5 μl 10% SDS and 100 μl 5 M potassium acetate. Proteins and tissue debris were pelleted by centrifugation at 3220 *g* for 30 min. The supernatant was saved and DNA was precipitated by adding 0.7 volume isopropanol and incubating at −20°C for at least 15 min. DNA was then pelleted by centrifugation at 3220 *g* for 45 min. DNA pellets were washed with 70% ethanol, briefly dried, and resuspended in 50 μl R25 (1×TE with 25 μg/ml RNase A). Typically, 2 μl was used in a 20 μl PCR reaction.

For *cref3* mutant genotyping, the following primer pairs were used.

For SALK_077977 T-DNA insertion,

LBb1.3: ATTTTGCCGATTTCGGAAC

SALK_077977_RP: ATCGAACACCTTACGTGCATC

For *CREF3* genomic DNA:

SALK_077977_LP: AAAGAGGATCTAACGGCGAAG
SALK_077977_RP: ATCGAACACCTTACGTGCATC

For *CREF3* transgene sequence verification, the following primer pairs were used. The forward primer sits in the *CREF3* presequence, and the reverse primer sits in the NOS terminator.

Forward: GGTCTCTCTAAATCAACCAAAC

Reverse: GCCAAATGTTTGAACGATCTGC

## Author contributions

YKS and IS designed the research. YKS performed the experiments. YKS, BG and IS collected, analysed and interpreted the data. YKS, BG and IS wrote the manuscript.

## Acknowledgements

The work was supported by grants from the Australian Research Council to IS (CE140100008 and FL140100179). YKS was recipient of a Research Training Program Scholarship from the Australian Government.

